# Deployment and transcriptional evaluation of nitisinone, an FDA-approved drug, to control bed bugs

**DOI:** 10.1101/2024.06.18.599347

**Authors:** M. Sterkel, J. Tompkin, C. Schal, L. M. Guerra, G. C. D. Pessoa, P. L. Oliveira, J. B. Benoit

**Affiliations:** Centro Regional de Estudios Genómicos, Facultad de Ciencias Exactas, Universidad Nacional de La Plata, CENEXA, CONICET, La Plata, Buenos Aires, Argentina; Department of Biological Sciences, University of Cincinnati, Cincinnati, OH 45221, USA; Department of Entomology and Plant Pathology, North Carolina State University, Raleigh, NC 27695, USA; Universidade Federal de Minas Gerais, Departamento de Parasitologia-ICB, Laboratório de Entomologia Médica, Belo Horizonte, MG, Brasil; Instituto de Bioquímica Médica Leopoldo de Meis, Universidade Federal do Rio de Janeiro, Rio de Janeiro, Brazil; Instituto Nacional de Ciência e Tecnologia em Entomologia Molecular, Rio de Janeiro, Brazil

**Keywords:** Hematophagous vector control. *Cimex lectularius*. Tyrosine catabolism. 4-hydroxyphenylpyruvate dioxygenase. Ectocide

## Abstract

Bed bugs are blood-feeders that rapidly proliferate into large indoor infestations. Their bites can cause allergies, secondary infections and psychological stress, among other problems. Although several tactics for their management have been used, bed bugs continue to spread worldwide wherever humans reside. This is mainly due to human-mediated transport and their high resistance to several classes of insecticides. New treatment options with novel modes of action are required for their control. In this study, we evaluated the use of nitisinone (NTBC), an FDA-approved drug, for bed bug control in an insecticide-susceptible (HH) and an insecticide-resistant (CIN) population. Although NTBC was lethal to both populations when administered orally or applied topically in very low doses, we observed a slight but significant resistance in the CIN population. Transcriptomic analysis in both populations indicated that NTBC treatment elicited a broad suppression of genes associated with RNA post-transcriptional modifications, translation, endomembrane system, protein post-translational modifications and protein folding. The CIN population exhibited higher ATP production and xenobiotic detoxification. Feeding studies on a mouse model highlight that NTBC could be used as a control method of bed bugs by host treatment. The results demonstrate that NTBC can be used as a new active ingredient for bed bug control by topical or oral treatment and shed light on the molecular mechanisms of suppressed tyrosine metabolism following NTBC treatment.

## 1. Introduction

The common bed bug, *Cimex lectularius* (Hemiptera: Cimicidae), is an indoor pest that poses serious global public health concerns by infesting not only residential settings but also hotels, cinemas, hospitals and public transport vehicles. Furthermore, bed bugs can be a significant pest of poultry facilities (González-Morales et al., 2022). Over the past two decades, bed bug infestations have resurged globally (Zorrilla-Vaca et al., 2015). Although bed bugs are not known to vector any human pathogens under natural conditions, their bites can cause secondary infections, allergies, psychological stress and other health issues (Akhoundi et al., 2023). Bed bug infestations are difficult to eliminate due to their rapid evolution of high resistance to commonly used insecticides such as pyrethroids, neonicotinoids and pyrroles. As a result, multiple treatments with different types of insecticides may be necessary to eradicate an infestation (Lee et al., 2023).

Various mechanisms of resistance have been described in bed bugs, including mutations in the target sites that reduce the binding of insecticides (Zhu et al., 2010), changes in the cuticle that decrease insecticide penetration (Lilly et al., 2016), and metabolic resistance that enhances the detoxification of insecticides (González-Morales et al., 2021). Specifically, the upregulation of certain detoxification enzymes like cytochrome P450-dependent monooxygenases (P450s), glutathione *S*-transferases (GSTs), carboxylesterases (CESTs), and esterases (ESTs) have been reported to be involved in metabolic resistance in different populations of bed bugs (Adelman et al., 2011; Bai et al., 2011; González-Morales et al., 2021). Other mechanisms such as sequestration of the insecticide, the microbiota, oxidative stress and avoidance behavioural adaptations have also been reported to influence insecticide resistance in other insects, such as mosquitoes (Ingham et al., 2023). Given the widespread resistance of bed bug populations to currently used insecticides and the pervasive cross-resistance to various insecticidal products, new active ingredients with different modes of action are needed to control bed bug populations.

Hematophagous arthropods can consume many times their body weight in a single blood meal, and they have evolved specific adaptations to prevent toxicity from the high quantities of amino acids, heme, iron, and salts generated during vertebrate blood digestion (Graça-Souza et al., 2006; Sterkel et al., 2017; Whiten et al., 2018). Specifically, it was recently shown that the tyrosine degradation pathway enables blood-feeding insects to tolerate high amounts of free tyrosine produced upon digestion of vertebrate blood proteins (Sterkel et al., 2016). Blocking this pathway causes the accumulation of toxic amounts of tyrosine during blood digestion, ultimately causing the death of blood-fed arthropods without affecting organisms that feed on other diets. This discovery led us to propose that inhibiting tyrosine catabolism might be a novel approach to selectively target hematophagous vectors (Sterkel et al., 2016). Inhibitors of the second enzyme in the tyrosine catabolism pathway, 4-hydroxyphenylpyruvate dioxygenase (HPPD), are commonly used as herbicides and in human health (Beaudegnies et al., 2009). Among the inhibitors tested for vector control, nitisinone (NTBC) was the most effective (McComic et al., 2023; Ramirez et al., 2021; Sterkel et al., 2021). NTBC is an orphan drug used to treat Tyrosinemia Type I (Lindstedt et al., 1992) and Alkaptonuria (Ranganath et al., 2016), rare medical conditions that require government intervention to subsidize drug development. When orally administered with the blood meal or topically applied, this drug was lethal to ticks, mosquitoes, tsetse flies, and kissing bugs (Cui et al., 2024; McComic et al., 2023; Ramirez et al., 2021; Sterkel et al., 2021, 2016).

In this study, we assessed the effectiveness of NTBC as a new method for controlling bed bugs in an insecticide-susceptible (Harold Harlan: HH) and an insecticide-resistant population (Cincinnati: CIN) through ingestion and topical application. The death of hematophagous arthropods is due to tyrosine accumulation (Sterkel et al., 2021, 2016), but the molecular and biochemical pathways affected by this drug are unknown. To gain insight into the effects of NTBC treatment on bed bug biology and the potential mechanisms of resistance to it, we performed a transcriptomic analysis in both populations. Altogether, the results suggest that NTBC could be an effective insecticide for controlling bed bugs by causing a general dysfunction in protein metabolism that can be used as an oral or topical treatment. This approach solely targets hematophagous ectoparasites and disease vectors, making it a more eco-friendly alternative to conventional insecticides.

## 2. Materials and methods

### 2.1. Ethics statement

Hairless mice obtained from the Universidade Federal de Minas Gerais (UFMG) were used for *in vivo* experiments. The animals were housed in standard plastic rodent cages with wood shavings as bedding material. The rodents were maintained on commercial food pellets, and water was provided *ad libitum*. These procedures and protocols were reviewed and approved by the Comissão de Ética no Uso de Animais (CEUA-UFMG 263/2021 – valid up to 3/27/2027).

### 2.2. Rearing of insects

The Harold Harlan (HH) population was collected in Ft. Dix, NJ, in 1973 and is also referred to as the Ft. Dix strain. They have been kept in plastic containers with cardboard shelters in an environmental chamber at 25°C, 50 ± 5% relative humidity and a 12-hour light-dark cycle at the University of Cincinnati (USA). This population has not been exposed to insecticides since 1973 and is therefore used as a reference strain for susceptibility testing (González-Morales et al., 2021). Similarly, the Cincinnati (CIN) population was collected in 2012 and reared in the same conditions as HH. Males from the CIN population have recently shown moderate resistance to fipronil and deltamethrin when applied topically (González-Morales et al., 2021). Both populations have been reared under laboratory conditions and fed on defibrinated rabbit blood (Hemostat, CA, USA) once a month with an artificial blood feeder (Hemotek, Blackburn, UK).

The bed bug population used in the *in vivo* studies was collected in 2014 and 2015 from public shelters in Belo Horizonte, Brazil (19° 55’ S 43° 56’ W). The insects were kept in a controlled environment at 27°C, 70% humidity, and 12 hours of light-dark cycles at the Federal University of Minas Gerais (Brazil) since that date, without the introduction of external material or exposure to insecticides.

### 2.3. Artificial feeding assay

Nitisinone (PHR1731, Sigma-Aldrich, St. Louis, MO, USA) was dissolved in PBS 1X (NaCl 137 mM, KCl 2.7 mM, Na2HPO4 10 mM and KH2PO4 1.8 mM) immediately before use. The pH was adjusted to 7-8 with 1M NaOH and 1M HCl. To achieve the final concentration for feeding, one volume of NTBC in PBS was mixed with nine volumes of defibrinated rabbit blood (Hemostat, CA, USA). Healthy insects, including first-instar nymphs, adult females and males of unknown age, were fed on rabbit blood mixed with NTBC through an artificial feeding apparatus (Hemotek) at 37 °C. Insects in the control group were fed on rabbit blood containing one volume of PBS 1X. The final NTBC concentrations in blood were 0.05, 0.1, 0.2, 0.3, 1, 3 and 9 µg/ml. The insects were held in the same conditions described above, and their mortality was assessed daily.

### 2.4. Topical application assay

Blood-fed adult insects received topical applications of NTBC dissolved in acetone using a 2 µl micropipette (Gilson) immediately after feeding. NTBC was dissolved in acetone immediately before use. Each bed bug received 0.5 µl of solution on the ventral thorax. The NTBC dosages applied were 750, 250, 83.3, 27.8, 9.3 and 3.1 ng/insect. Bed bugs in the control group received acetone only. The insects were kept in the same conditions described above and the mortality was assessed daily.

### 2.5. Ecdysis

The daily count of exuviae was used to determine the number of 1^st^ instar nymphs that moulted. Only insects that survived for more than five days after undergoing NTBC treatment were considered for calculating the percentage of successful ecdysis.

### 2.6. Oviposition and hatching assays

We used females of unknown ages, so all females were assumed to have mated before entering the assays. After undergoing NTBC treatment, the females were separated into individual vials and kept under the same conditions as before. Three weeks after the treatment, the number of eggs laid by each female was counted, along with the number of hatched nymphs. The ratio of hatching was calculated by dividing the number of hatched first-instar nymphs by the number of eggs laid by each female.

### 2.7. RNA isolation and RNA-Seq analysis

Thirty-five adult female insects were collected from the HH and CIN populations 24 hours after being fed on blood supplemented either with 0.3 µg/ml NTBC or PBS. Only live healthy insects were collected, and five insects were pooled in a 1.5 ml Eppendorf tube containing 1 ml of cooled TRIzol reagent (Invitrogen, CA, USA). Seven samples were collected for each condition. The insects were homogenized using BeadBlaster 24 microtube homogenizers (Benchmark Scientific, NJ, USA). RNA from the mixture was separated according to the TRIzol manufacturer’s protocol. Afterwards, DNA impurities were eliminated using DNase I (Thermo Scientific, PA, USA). The total RNA was then concentrated using the GeneJET RNA Cleanup and Concentration Micro Kit (Thermo Scientific, PA, USA). The concentration and quality of the RNA were checked using a Nanodrop spectrophotometer (Thermo Scientific, MA, USA). The samples were then sent in dry ice for sequencing to Novogene (Sacramento, CA, USA). The three samples that showed better RNA integrity number (RIN) values for each condition (HH-PBS, HH-NTBC, CIN-PBS and CIN-NTBC) were chosen for sequencing. The raw RNA-Seq data was uploaded to the National Center for Biotechnology Information (NCBI) Sequence Read Archive, Bio-project PRJNA1119180.

### 2.8. Quality assessment and analysis of RNA-Seq datasets

RNA-Seq analyses were performed as previously described (Benoit et al., 2023; Rosendale et al., 2022). Raw reads were trimmed using Trimmomatic (version 0.38) (Bolger et al., 2014) and the quality of reads was assessed with FastQC (version 0.11.5. https://www.bioinformatics.babraham.ac.uk/projects/fastqc/). Trimmed sequences were mapped to the bed bug genome GCA_000648675.3 (Benoit et al., 2016) using HISAT2 (version 2.1.0)(Kim et al., 2019). Gene count tables were generated with HTseq (https://htseq.readthedocs.io/en/latest/) under default settings.

Differentially expressed genes were determined using the generalized negative binominal model implemented in DESeq2 (version 1.18.1) in R (R Core Team, version 3.5.2). *P* values were corrected for multiple testing using the false discovery rate (FDR) approach and considered differentially expressed if the *P* value was smaller than or equal to 0.05. The analyses were conducted to compare the effect of NTBC treatment in the two populations (Cincinnati and Harlan) and a nested analysis was conducted to establish more general effects of NTBC treatment on bed bug biology for both HH and CIN populations. After the identification of the contigs with differential expression, enriched functional groups were identified with gProfiler (Raudvere et al., 2019) and clustered with REVIGO (Supek et al., 2011). The differentially expressed genes were searched against two reference protein databases (*Drosophila melanogaster* and SwissProt database) with the use of tBlastx. Functional annotations were assigned to each gene based on this comparison. The differentially expressed genes were categorized according to their known function or cell location in other insects, particularly *Drosophila melanogaster*.

### 2.9. *In vivo* feeding assay

Hairless mice were orally administered with different doses of NTBC dissolved in PBS 1X (0.5, 5, and 30 mg/kg body mass) while control mice received PBS 1X. Three replicates were performed for each dose using different mice on different days. The mice were anaesthetized intraperitoneally with 150 mg/kg ketamine and 10 mg/kg xylazine and ten adult females *C. lectularius* were allowed to feed on each treated mouse. In total, 30 insects were fed for each NTBC dose. The survival of the insects was monitored every 6 or 12 hours for 90 hours.

### 2.10. Statistical Analysis

For each dose in the artificial feeding and topical application assays, at least two independent experiments were conducted, with each experimental group consisting of 8-25 insects. The data from multiple experiments were combined to create a single graph. Statistical analysis and graph design were performed using Prism 8.0.2 software by GraphPad Software (San Diego, CA, USA). Survival analysis and the lethal time 50 (LT50: median time that it took 50% of insects to die) calculations were performed using Kaplan-Meier curves and Log-rank (Mantel-Cox) test analysis. On day ten after feeding, when the lethal doses reached 100% of mortality, the lethal concentration (Oral administration, LC50) or the lethal dose (Topical application, LD50) of NTBC that killed 50% of the insects of each population was determined using log-dose probit-mortality analysis in PoloPlus (LeOra Software Company, CA, USA). The toxicity of NTBC to the CIN population was compared to the HH population using a resistance ratio (RR50), which was calculated as CIN LD50/HH LD50. The confidence limits (CLs) of the RR50 were also calculated, and if the 95% confidence interval did not include the value of 1.0, then the LD50 values were considered statistically different (Wheeler et al., 2007).

To evaluate for significant differences in the ecdysis process, a two-way ANOVA (Dunnett’s multiple comparisons test) test was conducted to compare the groups fed on sublethal doses of NTBC (0.05, 0.1 and 0.2 µg/ml) and the group fed with PBS. To determine if there were any significant differences in the number of eggs laid by the females and their hatching between the NTBC-treated and control groups, a one-way ANOVA (Dunnett’s multiple comparisons test) was conducted.

## 3. Results

### 3.1 Survival analysis of *C. lectularius* following NTBC oral administration through artificial feeding

Artificial feeding assays were conducted with first-instar nymphs and adult males and females of HH and CIN bed bug populations using defibrinated rabbit blood supplemented with different concentrations of NTBC (ranging from 0.05 to 9 µg/ml). The LC50 values (concentration that killed 50% of the insects 10 days after feeding) for nymphs were 0.17 and 0.26 µg/ml for the HH and CIN populations, respectively (Fig. 1A-C). According to (Sierras and Schal, 2017), first-instar bed bug nymphs consume approximately 0.5 µl of blood; therefore, the ingested LD50 HH and CIN nymphs were around 0.08 ng and 0.12 ng of NTBC, respectively. The estimated RR50 was 1.52 (95% CI=1.01-2.11), indicating a slight but statistically significant resistance in CIN nymphs (Fig. 1A-C). The LT50 for lethal doses, which killed all bed bugs within 10 days, was on average 4 days PBM (post-blood meal; Fig. S1A-B, Table 1). The administration of NTBC did not affect the ecdysis process and almost all of the nymphs that survived longer than 5 days PBM moulted to the next instar (Fig. S1C).

**Figure 1:**
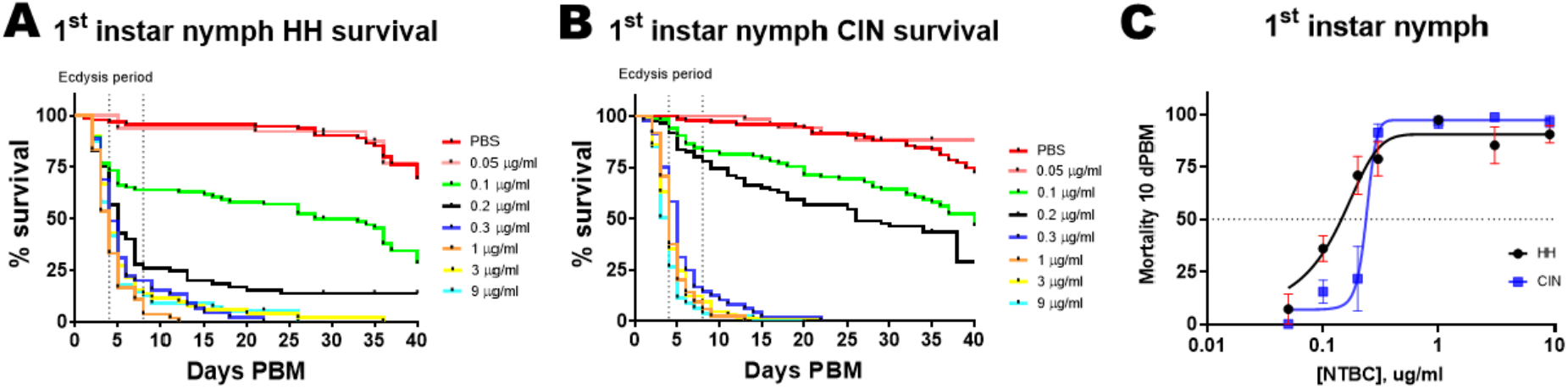
Oral administration of NTBC is lethal to *Cimex lectularius* nymphs. **A)** Survival of the Harold Harlan (HH) and **B)** Cincinnati (CIN) populations fed rabbit blood supplemented with NTBC or PBS. Three to five independent replicates were performed for each dose. The data from all replicates within a dose were combined for survival analysis and are represented by a single line in each graph. The total number of HH and CIN nymphs used was 513 and 587, respectively. **C)** Dose-response curves are shown for HH and CIN nymphs 10 days PBM (postblood meal), with each point representing mean±SEM. All NTBC doses, except 0.05 µg/ml, resulted in higher mortality than controls.

**Table 1.** Lethal time 50 (Days) for the different doses in artificial feeding assays.

|  | PBS | 0.1 µg/ml | 0.2 µg/ml | 0.3 µg/ml | 1 µg/ml | 3 µg/ml | 9 µg/ml | LC<sub>50</sub> 10 dPBM (µg/ml) | RR<sub>50</sub> (CIN/HH) | 95% limits |
| --- | --- | --- | --- | --- | --- | --- | --- | --- | --- | --- |
| N1 HH | 40.7 | 19.8 | 4.33 | 5.5 | 3.8 | 4.75 | 3.9 | 0.171 | 1.52 | 1.10 to 2.10 |
| N1 CIN | 45.1 | 22.1 | 23.0 | 4.4 | 4.0 | 3.7 | 3.4 | 0.26 |  |  |
| Female HH | 41.67 | 26.0 | 9.5 | 2.5 | 2.5 | 3.0 | 2.5 | 0.151 | 1.64 | 1.11 to 2.43 |
| Female CIN | 47.0 | 31.17 | 30.0 | 2.75 | 2.0 | 2.5 | 3.0 | 0.248 |  |  |
| Male HH | 44.75 | 35.0 | 23.0 | 6.5 | 9.0 | 9.25 | 6.75 | 0.414 | 1.18 | 0.51 to 2.46 |
| Male CIN | 39.75 | 32.33 | 27.0 | 8.5 | 8.25 | 6.75 | 8.5 | 0.489 |  |  |

The oral administration of NTBC to adult insects was also lethal. The LC50 for females in the HH and CIN populations was 0.15 and 0.25 µg/ml, respectively, with an estimated RR50 of 1.64 (95% CI=1.11-2.43; Fig. 2 A-C.). Assuming an ingested blood volume of approximately 3.92 µL (Sierras and Schal, 2017), the oral NTBC LD50 values were 0.59 and 0.97 ng, respectively. The LT50 was 2.6 days (Fig. S2A-B, Table 1), and 100% of mortality was achieved 5 days after feeding with lethal doses. The oral administration of sublethal doses of NTBC did not affect the reproductive fitness of females. There were no differences in the number of eggs laid (Fig. S2C) or their hatch rate (Fig. S2F) between NTBC-treated females and controls.

**Figure 2:**
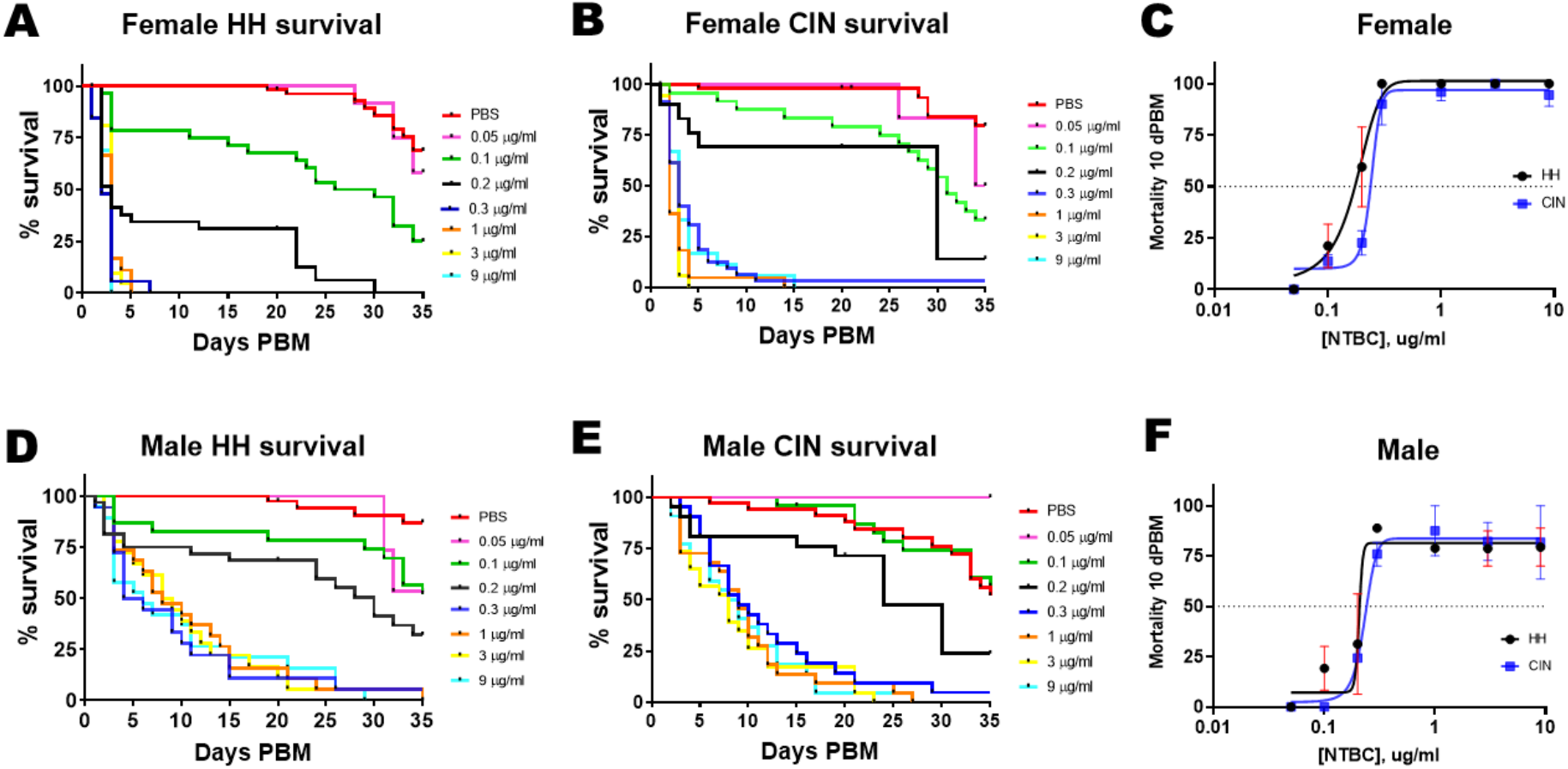
Oral administration of NTBC is lethal for adult *Cimex lectularius*. Survival of the Harold Harlan (HH) **A)** females and **D)** males, and Cincinnati (CIN) **B)** females and **E)** males fed rabbit blood supplemented with NTBC or PBS. Two independent replicates were performed for each dose. The data from the two independent replicates for each treatment were combined for survival analysis and represent a single line in each graph. We used 182 HH and 149 CIN females, and 184 HH and 167 CIN males. Dose-response curves are shown for **C)** females and **F)** males 10 days PBM (post-blood meal), with each point representing mean±SEM. All NTBC doses, except 0.05 µg/ml, resulted in higher mortality than controls. In CIN males, the lethality of 0.1 µg/ml was not statically different from controls.

The mortality rate of males showed a different pattern. After feeding, it took them a longer time to die, with an LT50 of around 9 days (Fig. S2D-E, Table 1), and 100% of mortality was achieved 25 days after feeding on lethal doses (Fig. 2D-E). On day 10 PBM, the LC50 values were found to be 0.414 and 0.489 µg/ml for HH and CIN, respectively (Fig. 2F). Assuming a blood volume ingested of approximately 3.92 µL (Sierras and Schal, 2017), the oral NTBC LD50 values were estimated to be 1.6 ng and 1.9 ng, respectively. The RR50 was 1.18 (with 95% CI=0.51-2.46), and no significant differences were observed in the susceptibility to NTBC between HH and CIN males when it was orally administered.

### 3.2. Survival analysis of *C. lectularius* following NTBC topical application

Since most bed bug control strategies nowadays rely on insecticide spray applications that penetrate through the bugs’ cuticles, we determined the effectiveness of NTBC through topical applications. Results indicate that the LD50 for HH and CIN females was 20 and 32.5 ng/female, respectively. The RR50 was 1.64 (95% CI=1.01-2.65), similar to that observed for the artificial feeding assays, suggesting that the cuticle might not be involved in resistance towards NTBC in CIN females and pointing to metabolic resistance (Fig. 3A-C, Table 2). The data indicate that when administered orally to females from both populations, NTBC was around 33 times more potent than when topically applied (Table 3). The LT50 for lethal doses was on average 2.7 days (Fig. S3A-B. Table 2). As observed in artificial feeding assays, sublethal doses of NTBC topically applied did not affect the reproductive fitness of females (Fig. S3C, F).

**Figure 3.**
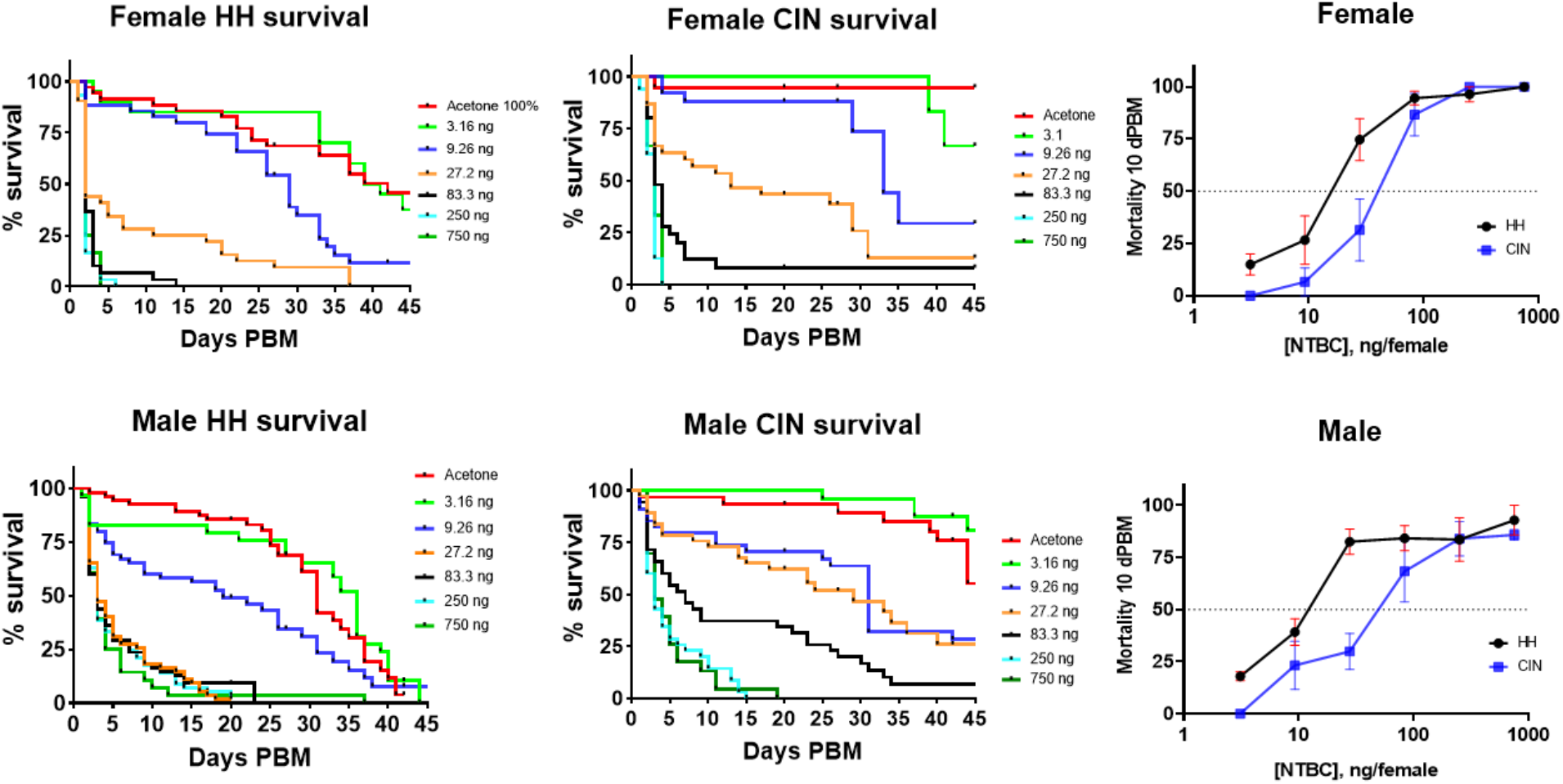
Topical application of NTBC is lethal for adult *Cimex lectularius*. Survival of the Harold Harlan (HH) **A)** females and **D)** males, and Cincinnati (CIN) **B)** females and **E)** males, topically treated with NTBC or acetone as control (0.5 µl/insect). Two to four independent replicates were performed for the different doses. The data from the independent replicates for each treatment were combined for survival analysis and represent a single line in each graph. We used 244 HH and 124 CIN females and 334 HH and 221 CIN males. Dose-response curves are shown for **C)** females and **F)** males 10 days PBM (post-blood meal), with each point representing mean±SEM. All NTBC doses, except 3.1 ng/insect, resulted in higher mortality than controls.

**Table 2.** Lethal time 50 (Days) for the different doses in topical application assays.

|  | Acetone 100% | 3.1 ng | 9.26 ng | 27.2 ng | 83.3 ng | 250 ng | 750 ng | LD<sub>50</sub> 10 dPBM (ng) | RR<sub>50</sub> (CIN/HH) | 95% limits |
| --- | --- | --- | --- | --- | --- | --- | --- | --- | --- | --- |
| Female HH | 43.0 | 42.0 | 22.25 | 3.5 | 2.37 | 2.25 | 2.0 | 19.9 | 1.64 | 1.01 to 2.65 |
| Female CIN | >50 | >50 | 34.0 | 16.5 | 4.17 | 2.5 | 3.0 | 32.5 |  |  |
| Male HH | 32.5 | 33.0 | 17.17 | 4.37 | 4.37 | 4.25 | 3.0 | 14.3 | 4.33 | 2.23 to 8.3 |
| Male CIN | 47.5 | >50 | 24.67 | 17.0 | 9.1 | 3.1 | 3.5 | 61.9 |  |  |

**Table 3:** LC50 and LD50 for artificial feeding assays calculated based on the volume of blood ingested (Sierras and Schal, 2017), and the LD50 calculated on the topical application assays. The last column shows the ratio between LD topical/LD oral.

|  | Artificial feeding | Artificial feeding | Topical application |  |
| --- | --- | --- | --- | --- |
|  | LC<sub>50</sub> 10 dPBM (µl/ml) | LD<sub>50</sub> 10 dPBM (ng/insects) | LD<sub>50</sub> 10 dPBM (ng/insect) | LD topical/LD oral |
| N1 HH | 0.17 | 0.08 | - |  |
| N1 CIN | 0.26 | 0.12 | - |  |
| Female HH | 0.15 | 0.59 | 19.9 | 33.82 |
| Female CIN | 0.25 | 0.98 | 32.56 | 33.22 |
| Male HH | 0.41 | 1.61 | 14.3 | 8.9 |
| Male CIN | 0.49 | 1.92 | 61.9 | 32.23 |

When NTBC was topically applied to males, moderate resistance was observed in the CIN population. The LD50 values on day 10 PBM were 14.3 and 61.9 ng/male for HH and CIN, respectively, and the LT50 for lethal doses was 4.5 days (Fig S2D-E). The RR50 was 4.3 (95% CI=2.23-8.3; Fig. 3D-F, Table 2). This is different from the results obtained in the artificial feeding assays, where no resistance was observed in CIN males and the LT50 was 9 days. However, a delay in the rate of death of males in comparison to females was observed, as with artificial feeding assays. These findings indicate that the cuticle plays a role in the resistance of CIN males to NTBC. When given orally, NTBC was approximately 9 and 32 times more potent for HH and CIN males, respectively (Table 3).

### 3.3. Transcriptomic analysis on HH and CIN females fed NTBC

#### 3.3.1. General description of transcriptional changes

The molecular pathways and mechanisms leading to death of blood-feeding arthropods due to tyrosine accumulation after NTBC treatment are still unknown. Besides, it is important to understand the mechanisms that might provide resistance to NTBC. To gain insight into how bed bugs respond to NTBC treatments, we conducted a transcriptomic analysis on HH and CIN females fed blood supplemented with PBS only (control group) or NTBC in PBS that resulted in 0.3 µg/ml blood (the lower lethal dose for both populations). We analyzed the transcript log2 (fold change) values 24 hours after feeding. The analysis revealed significant differential expression of many transcripts; 86 and 61 genes were downregulated in HH and CIN females, respectively; 19 of them were downregulated in both populations. The upregulated genes were 50 and 55 in HH and CIN respectively; 8 of them were upregulated in both populations (Fig. S4). GO term analysis showed that the downregulated transcripts are associated with the general processes of protein metabolism, which includes protein folding (GO: 0051082), translational initiation (GO:0006413), and ncRNA metabolic processes (GO:0034660) (Fig. 4). The transcripts upregulated in NTBC-treated bed bugs resulted in enriched GO categories for the HH population that included factors associated with aspartic-type peptidase activity (GO:0004190). In contrast, the major enriched categories upregulated in the CIN population include ATP synthesis (Fig. 4). A nested analysis with both populations indicates that the nucleoside triphosphate biosynthetic process (GO:0009141) is generally increased in both populations (Fig. 4).

**Figure 4.**
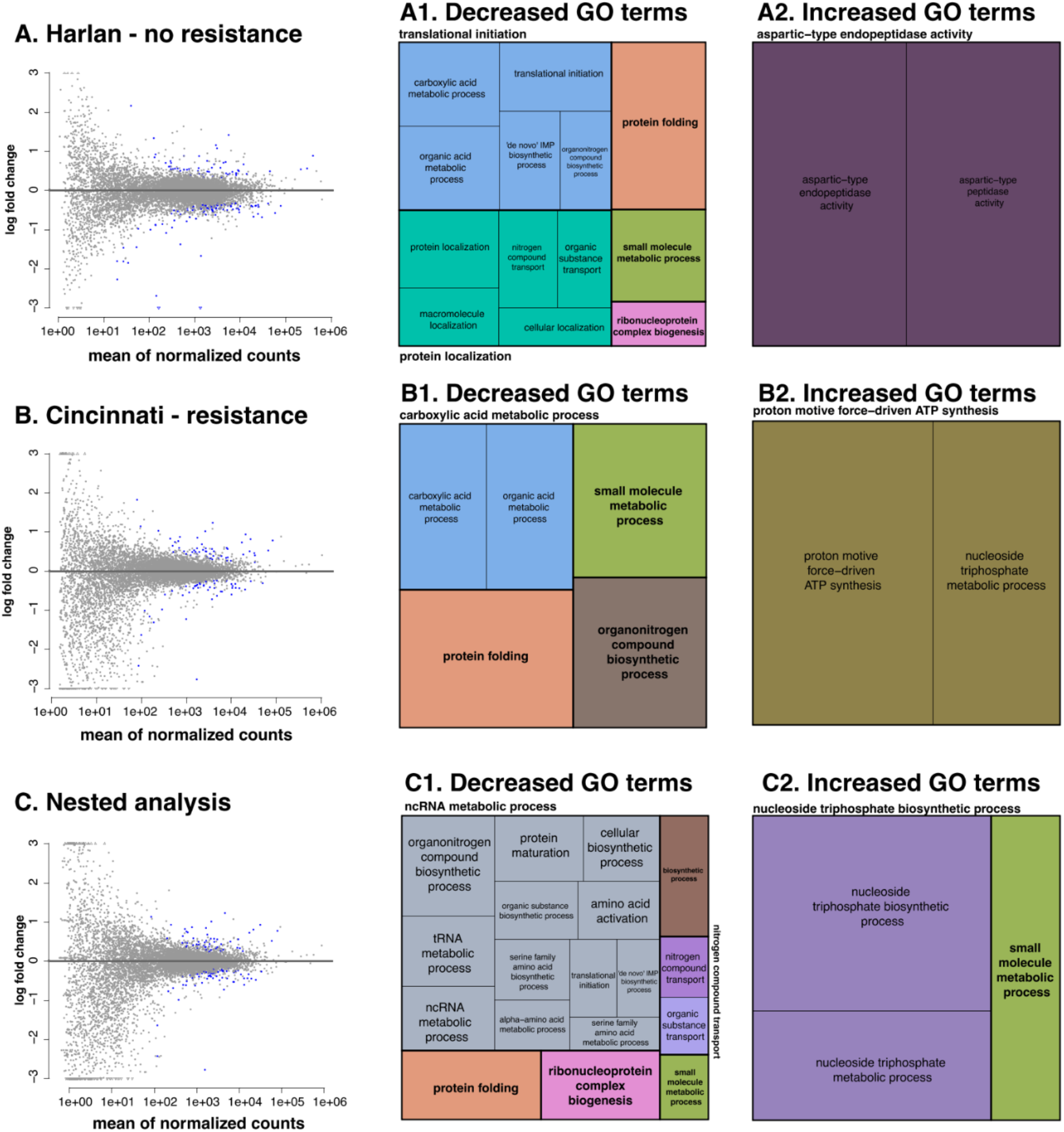
Dispersion plots and gene ontologies (GO) of interest and their relevant molecular functions, cellular components, and biological processes were visualized using treemaps in NTBC-treated **A)** HH and **B)** CIN females. **C)** Nested analyses considering both populations. Blue dots are genes with significant differential expression (FDR, *P* < 0.05). The boxes in the treemaps represent the unique functional categories while the colours represent the major GO groups. The treemaps were generated using REVIGO.

#### 3.3.2. Targeted description of transcriptional changes associated with gene groups of interest

This analysis revealed changes in mRNA levels of genes that encode proteins regulating RNA polymerase II transcriptional activity in both populations (Table S1), suggesting that the activity of this polymerase may be altered. The analysis also revealed the downregulation of many transcripts encoding proteins associated with RNAs (mRNA, rRNA and tRNA) post-transcriptional modifications and ribosome biogenesis (Table S2). In addition, many subunits of different eukaryotic translation initiation factors (eIF2, eIF3 and eIF4) were downregulated (Table S3). Moreover, many transcripts coding proteins that play a role in the endomembrane system were also downregulated in both populations, including proteins involved in protein transport and post-translational modifications (Table S4). Additionally, numerous chaperones and proteasomal proteins were downregulated (Tables S5, S6). Peptidase activity was also altered (Table S7).

We also observed that the mRNA levels of many transmembrane transporters (organic and inorganic) were altered. Many of them are mainly expressed in the central nervous system, suggesting that NTBC treatment and/or tyrosine accumulation may impact neural functions (Table S8). A downregulation in sterol transporters (Niemann-Pick type protein-2 (NPC-2) and NPC-1B) was also observed in both populations (Table S9).

The mRNA levels of genes encoding enzymes for various metabolic pathways were changed. ATP-citrate synthase and glutamine synthetase were downregulated in both populations (Table S10). Interestingly, tyrosine aminotransferase (TAT), the first enzyme in the tyrosine catabolism pathway, was among the genes that were more upregulated in both populations. This indicates that the bed bugs respond to tyrosine accumulation due to the inhibition of HPPD by increasing TAT transcription. Additionally, 4-coumarate-CoA ligase, an enzyme involved in the phenylpropanoid biosynthetic pathway, was upregulated in both populations (Table S10).

The ATP synthase subunits beta, gamma, delta, and epsilon, along with other proteins related to mitochondria function, were up-regulated exclusively in the insecticide-resistant CIN population. Two peroxiredoxins, which play a role in antioxidant function in the mitochondria, were also upregulated (Table S11). Furthermore, four genes coding for enzymes associated with xenobiotic detoxification were upregulated only in this population. These enzymes include two glutathione *S*-transferases (Sigma-1 and theta-1), a carboxylesterase, and a cytochrome P450 9e2 (Table S12).

Finally, many uncharacterized genes presented altered mRNA levels upon NTBC treatment. The most downregulated gene in both populations is uncharacterized (LOC106661977), and its function has not been described in any organism. A leucine-rich repeat-containing protein 4 (LOC106665058) was also downregulated in both populations. Besides, four upregulated genes in both populations are uncharacterized (LOC106674390, LOC106674306, LOC106661024, LOC106670295), two of them are described as “probable salivary secreted peptid” (Table S13).

### 3.4. Direct evaluation of host treatment with NTBC for control of *C. lectularius*

People with Tyrosinemia Type I consume a therapeutic oral dose of 1 mg NTBC/kg/day, which results in an NTBC plasma concentration of 8.2 µg/ml, and has a half-life of 54 hours (Hall et al., 2001). The results of artificial feeding assays suggested that NTBC could be administered to vertebrate hosts as an ectocide to control bed bugs. To test this hypothesis, we conducted *in vivo* experiments. Hairless mice were orally administered with NTBC in three doses: 0.5, 5 and 30 mg/kg, and *C. lectularius* females were fed on them 2-3 hours after drug administration. All doses resulted in lethality, with LT50 around 2 days PBM (Fig. 5), similar to the LT50 observed in the artificial feeding assays. This provides direct evidence that NTBC can be used as an oral treatment for bed bugs control.

**Figure 5:**
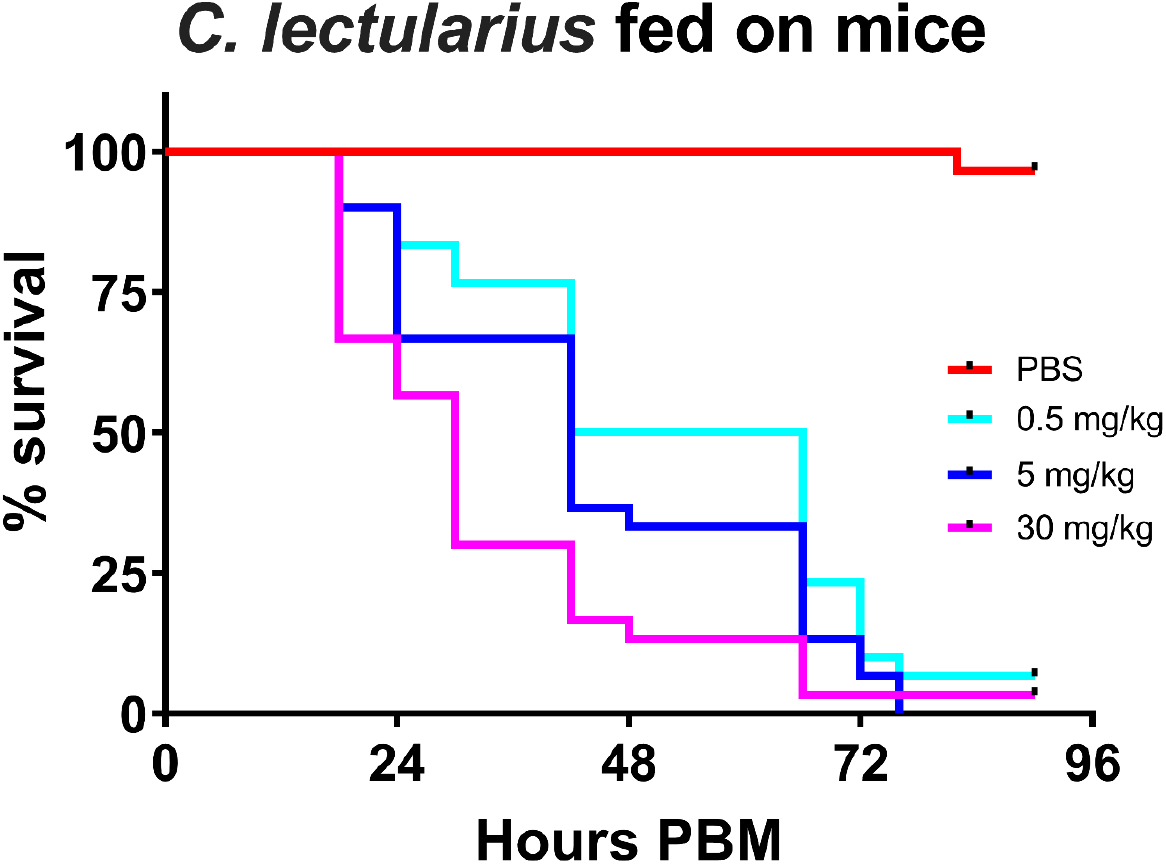
Feeding on mice treated with an oral dose of NTBC is lethal for *C. lectularius*. Three mice were treated with each dose (twelve animals in total) and 10 adult female bed bugs were fed on each mouse. The total number of bed bugs used was 120. Data are shown as Kaplan-Meier curves. All doses resulted in higher mortality than controls. This experiment used insects obtained from a *C. lectularius* colony established in the Universidade Federal de Minas Gerais from field-collected bed bugs.

## 4. Discussion

The primary method for mitigating indoor bed bug infestations involves using insecticides. However, due to the global increase in bed bug populations and their resistance to currently used pesticides (Lee et al., 2023), new alternatives are urgently needed. Our study indicates that inhibiting tyrosine catabolism with NTBC could be a novel approach for controlling bed bugs. Bed bugs are more susceptible to NTBC compared to other blood-feeding insects such as kissing bugs (Sterkel et al., 2016), mosquitoes (McComic et al., 2023; Sterkel et al., 2016; Vergaray Ramirez et al., 2022), tsetse flies (Sterkel et al., 2021) and ticks (Cui et al., 2024). The insects die due to the buildup of tyrosine from digested proteins, and the time it takes for them to die depends on how quickly they digest their blood meal (Sterkel et al., 2016). We observed unexpected differences in the response to NTBC based on the sex of the bed bugs. Males showed a higher LT50 in both insecticide-susceptible and resistant populations. This difference could be due to a slower digestion rate in males or other sex-related differences in bed bugs. Further investigation is needed to understand the reasons behind these differences.

A small but statistically significant resistance was estimated for CIN nymphs and females (RR50=1.6) upon oral administration, while no resistance was observed in males. Furthermore, the LC for males in both populations was approximately twice as high as in females from the respective populations. Since the LC50s were calculated on day 10 after feeding for both males and females, the lower rate of male death increases their LCs and may mask the small resistance that we observed in females. Indeed, the dose of 0.1 µg/ml caused higher mortality in HH males than in the HH controls, but was not different in CIN males from the CIN controls, indicating a difference between HH and CIN male susceptibility to NTBC despite not being statistically significant.

As observed when NTBC was orally administered to bed bugs, differences related to the insects’ sex were also observed upon NTBC topical application, with a delay in mortality in males compared with females. The RR50 for females in topical application assays was similar to that calculated when NTBC was orally administered. However, different from artificial feeding assays, resistance was observed in CIN males (RR50=4.3). The potency of NTBC was similar to that reported for fipronil when topically applied to HH and CIN males (LD50 values equal to 20.3 ng and 167 ng, respectively (RR50=8.4)). The CIN males were also resistant to deltamethrin. In this population, P450s were associated with fipronil resistance (González-Morales et al., 2021). Our data suggest that NTBC resistance in CIN males may be associated with the cuticle since no differences were observed in males when NTBC was orally administered. Nevertheless, given the small RR50 values, two important caveats might apply to all our oral and topical assays. First, differences in body mass between the two populations and between females and males, if any, might require LD50 and LT50 values to be normalized per mg or gram body mass. Secondly, differences in the amount of blood ingested by the two populations and between females and males might also affect the estimated toxicity values and slightly reduce or increase the RR50 values.

The administration of sublethal doses of NTBC did not affect the reproductive fitness of females. Besides, females treated with lethal doses laid some eggs before they died. This fact is important in preventing the fixation of resistant alleles in the population, as it reduces the selective pressure, similar to the idealized late-acting insecticides proposed for malaria control (Koella et al., 2009; Read et al., 2009). This particular characteristic of NTBC is different from neurotoxic insecticides that kill immediately after application, preventing reproduction.

Although tyrosine accumulation and precipitation have been proposed as the mechanism responsible for hematophagous arthropod death after a blood meal due to HPPD inhibition (Sterkel et al., 2021, 2016), little is known about the molecular mechanisms affected by NTBC and tyrosine. In this study, we identified several cellular processes affected in HH and CIN populations. The RNA-Seq results revealed downregulation of transcript levels of several genes associated with RNA post-transcriptional modifications, translation, endomembrane system transport, post-translational protein modifications and folding, and protein degradation. In hematophagous arthropods, an increase in protein synthesis occurs coupled with blood meal digestion (Lehane, 2005), the downregulation of these processes in NTBC-treated insects may indicate that the increase in protein synthesis that normally occurs after a blood meal does not take place.

The levels of many organic and inorganic membrane transporters, some of which are located in the nervous system, were found to be disturbed in both populations. In humans, mental disorders are observed in the different tyrosinemias (Scott, 2006). Further evaluation is needed to understand the impact of NTBC and tyrosine accumulation on the nervous system of blood-feeding insects and its association with the lethal phenotype.

Importantly, we identify distinct pathways upregulated in the insecticide-resistant CIN females upon NTBC ingestion, including mitochondrial ATP production and xenobiotic detoxification. The increased peroxidase mRNA levels observed are likely a response to the increased mitochondrial respiration and the consequent production of H2O2. Oxidative stress plays a role in or is a result of resistance to pyrethroids in anopheline mosquitoes. Changes in the redox state have been observed post-pyrethroid exposure (Ingham et al., 2021; Ingham et al., 2023). Besides, an increase in the expression of genes within the oxidative phosphorylation pathway was observed in two *Anopheles coluzzii* resistant populations compared to the susceptible control, which translated phenotypically through an increased respiratory rate (Ingham et al., 2021). The increased mitochondrial activity observed in the CIN population upon NTBC ingestion would help counter the effects of the drug and may contribute to resistance.

Previously, transcriptomic analysis performed in bed bugs showed that strains resistant to deltamethrin presented upregulated transcript levels of cytochrome P450 and carboxylesterase (Adelman et al., 2011). Also, a comparison of transcriptomes of pesticide-resistant and pesticide-susceptible bed bugs indicated the upregulation of transcripts involved in penetration resistance and metabolic resistance (Mamidala et al., 2012). Our transcriptional analysis identified four detoxification genes (two GSTs, one P450 and one CEST) that were upregulated in CIN females fed on NTBC. This upregulation was not observed in HH females. These enzymes could be involved in the small but significant resistance observed in the CIN population towards NTBC, and perhaps the possible cross-resistance observed with fipronil and deltamethrin in this population.

The effectiveness of certain insecticides was tested for their potential use in creating a liquid bait for bed bugs. Abamectin, clothianidin, and fipronil caused 100% mortality by day 3, while indoxacarb and its bioactive metabolite DCJW were ineffective. Fipronil was the most potent, presenting an LC50 in HH adult males equal to 13.4 ng/ml blood (0.052 ng/male) (Sierras and Schal, 2017). The data presented here indicate the potential use of NTBC in bed bug baits. Moreover, *in vivo* experiments demonstrated that doses as low as 0.5 mg/kg resulted in almost 100% bed bug mortality 3-4 days after they fed on mice treated with NTBC, confirming that it could be used as an ectocide to control bed bugs. The pharmacokinetic profile in humans (Hall et al., 2001) indicates that the administration of a single therapeutic dose (1 mg/kg) would maintain NTBC blood concentrations above the bed bug LC50 (0.25 µg/ml) for around 11 days. Therefore, weekly dosing with NTBC would indefinitely extend its lethal effects on bed bugs. Systemic veterinary drugs are commonly used to control ectoparasites in companion animals and livestock. These drugs have been proposed for ectoparasite control in humans (Miglianico et al., 2018), but with a few exceptions (e.g., ivermectin), their pharmacokinetic properties and potential side effects have not been thoroughly assessed. As bed bugs also proliferate in poultry farms and impact the health and welfare of chickens and workers, systemic drugs like the isoxazoline fluralaner (González-Morales et al., 2023, 2022), and possibly NTBC, have great potential in ectoparasite control. These novel treatments could be used individually or combined with other pesticides to be more effective control treatments. Since NTBC has been used in medicine for more than 30 years, it presents the advantage that it could be administered also to humans.

Collectively, our study provides a foundation for the possible use of NTBC for bed bug control and sheds light on the molecular processes affected by this drug. Moreover, this study may help to elucidate the mechanisms responsible for the low level of NTBC resistance we detected in the CIN population. These mechanisms should be further studied by performing functional assays, such as RNA interference experiments targeting the putative candidate genes and analyzing the susceptibility of bed bugs toward NTBC. This would help devise future strategies to control bed bug populations and manage resistance.

## 5. Conclusions

NTBC was highly effective at very low concentrations when given orally and applied topically to bed bugs. The low level of resistance in one field-collected population needs to be assessed with more populations because it falls within the range of variation of such assays. RNA-Seq analyses revealed potential mechanisms of NTBC action, which are likely related to general dysfunction of translation and protein processing. Additionally, oral administration of a low dose of NTBC to a vertebrate host killed nearly 100% of bed bugs colonized from a field collection about a decade ago. These results suggest that NTBC administered orally to vertebrate hosts, including humans, could be a viable method for controlling bed bugs. If used as part of an integrated management strategy, NTBC could improve the elimination of bed bugs. Furthermore, repurposing NTBC, an FDA-approved drug, for bed bug control may help overcome many regulatory hurdles.

## Acknowledgements

We are grateful to Fulbright (Argentina-United States) and CONICET (Argentina) for providing a fellowship that allowed us to carry out this work. We would also like to express our gratitude to Rick Santangelo and Melissa Uhran for raising the colonies of HH and CIN bed bugs. Partial funding for shared equipment was provided by the National Institute of Allergy and Infectious Diseases under Award Numbers R01AI148551, R21AI166633, and R21AI176098 (JBB).

