## Supplemental Figures for "Deployment and transcriptional evaluation of nitisinone, an FDA-approved drug, to control bed bugs"

**Supplemental data**

**Supplemental figures**

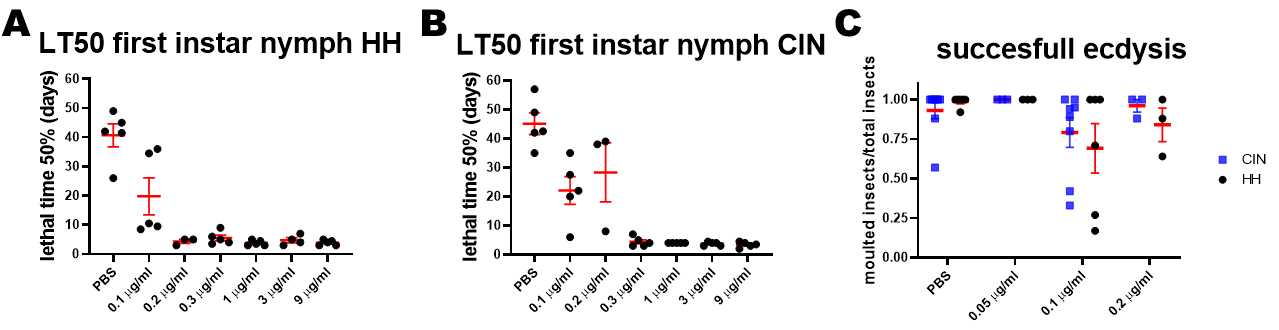

**Figure S1.** Lethal time 50 (LT_50_, time that takes 50% of the insects to die) for the different NTBC doses in *C. lectularius* first instar nymphs **A)** Harold Harlan (HH) and **B)** Cincinnati (CIN) populations. **C)** Percentage of successful ecdysis after feeding on different doses of NTBC. The percentage of successful ecdysis was calculated as (nymphs that survived longer than 5 days after feeding/number of moulted nymphs) for each independent experiment. The ecdysis period was between four to eight days after feeding. Each point represents an independent experiment. Data are shown as mean±SEM.

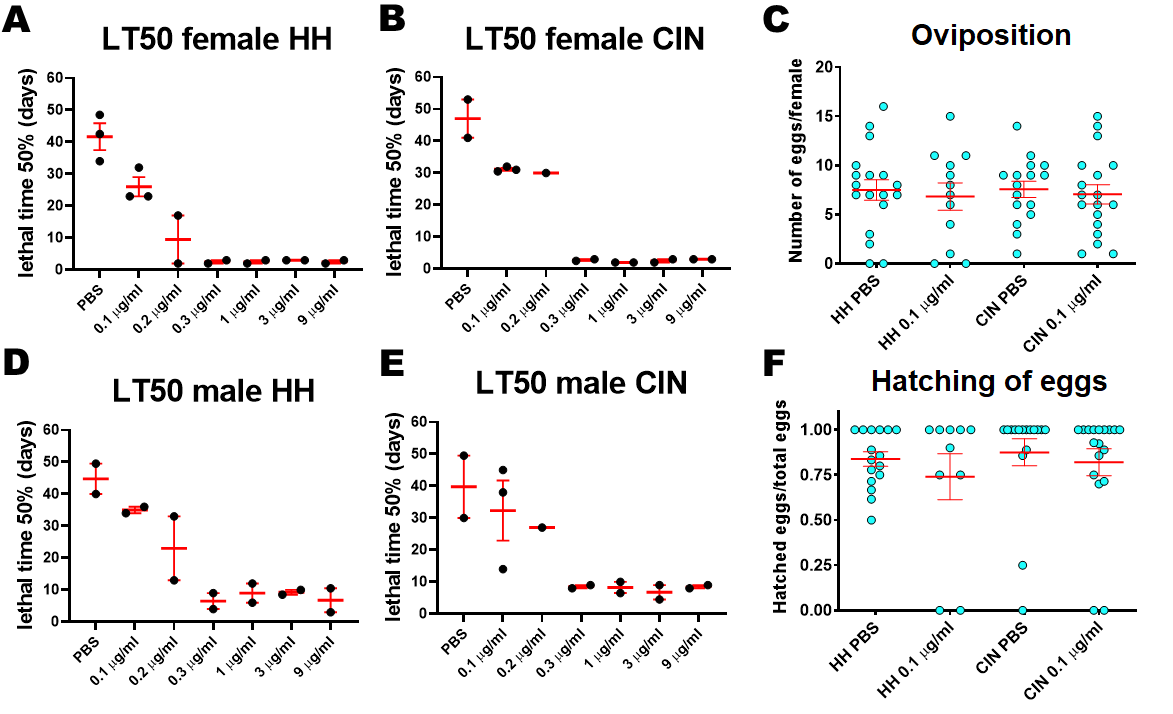

**Figure S2.** Lethal time 50 (LT_50_, time that takes 50% of the insects to die) in artificial feeding assays for the different NTBC doses in *C. lectularius* females **A)** Harold Harlan (HH) and **B)** Cincinnati (CIN) populations. **C)** Number of eggs laid by each female fed with a sub-lethal NTBC dose. LT_50_ in artificial feeding assays for different NTBC doses in **D)** HH males and **E)** CIN males. **F)** Hatching of eggs calculated as (first instar nymphs hatched/eggs laid by each female). Data are shown as mean±SEM.

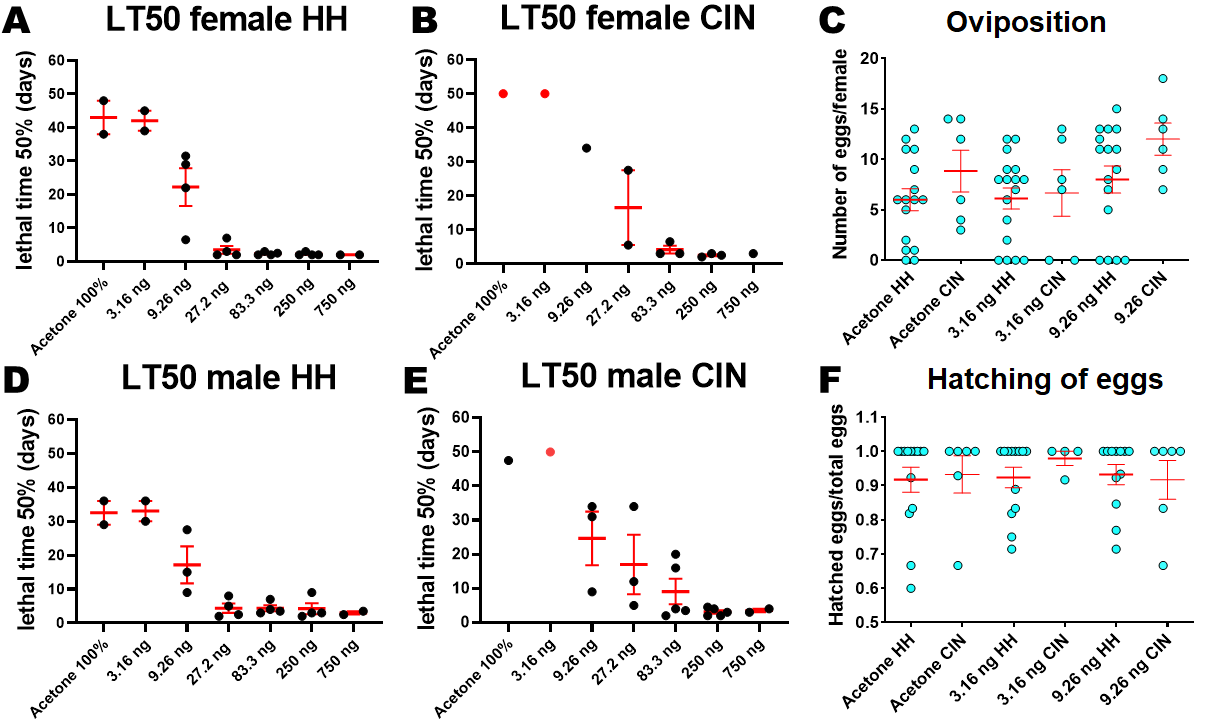

**Figure S3.** Lethal time 50 (LT_50_, time that takes 50% of the insects to die) in topical application assays for the different NTBC doses in *C. lectularius* females **A)** Harold Harlan (HH) and **B)** Cincinnati (CIN) populations. **C)** Number of eggs laid by each female fed with a sub-lethal NTBC dose. LT50 in topical application assays for different NTBC doses in **D)** HH *C. lectularius* males and **E)** CIN males. **F)** Hatching of eggs calculated as (first instar nymphs hatched/eggs laid by each female). Data are shown as mean±SEM. The red points in **B)** and **E)** indicate that the LT_50_ values were higher than 50 days, when the experiments ended.

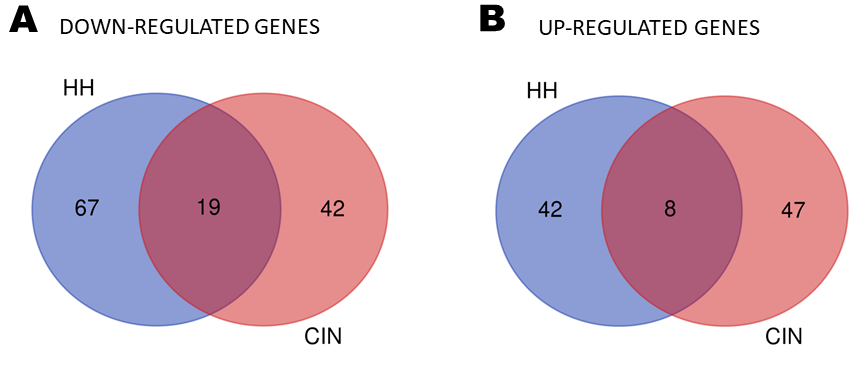

**Figure S4.** The number of genes that presented altered mRNA levels after 0.3 µg/ml NTBC treatment, compared to controls (PBS treatment), in HH and CIN females. **A)** Downregulated genes. **B)** Upregulated genes. Venn diagrams were generated using Bioinformatics & Evolutionary genomics. (http://bioinformatics.psb.ugent.be/webtools/Venn/).
